# ERNIE-RNA: An RNA Language Model with Structure-enhanced Representations

**DOI:** 10.1101/2024.03.17.585376

**Authors:** Weijie Yin, Zhaoyu Zhang, Liang He, Rui Jiang, Shuo Zhang, Gan Liu, Xuegong Zhang, Tao Qin, Zhen Xie

**Author notes:** Equal first authorship (Weijie Yin, Zhaoyu Zhang, Liang He).

## Abstract

With large amounts of unlabeled RNA sequences data produced by high-throughput sequencing technologies, pre-trained RNA language models have been developed to estimate semantic space of RNA molecules, which facilities the understanding of grammar of RNA language. However, existing RNA language models overlook the impact of structure when modeling the RNA semantic space, resulting in incomplete feature extraction and suboptimal performance across various downstream tasks. In this study, we developed a RNA pre-trained language model named ERNIE-RNA (**E**nhanced **R**eprese**n**tations with base-pa**i**ring r**e**striction for **RNA** modeling) based on a modified BERT (Bidirectional Encoder Representations from Transformers) by incorporating base-pairing restriction with no MSA (Multiple Sequence Alignment) information. We found that the attention maps from ERNIE-RNA with no fine-tuning are able to capture RNA structure in the zero-shot experiment more precisely than conventional methods such as fine-tuned RNAfold and RNAstructure, suggesting that the ERNIE-RNA can provide comprehensive RNA structural representations. Furthermore, ERNIE-RNA achieved SOTA (state-of-the-art) performance after fine-tuning for various downstream tasks, including RNA structural and functional predictions. In summary, our ERNIE-RNA model provides general features which can be widely and effectively applied in various subsequent research tasks. Our results indicate that introducing key knowledge-based prior information in the BERT framework may be a useful strategy to enhance the performance of other language models.

## Introduction

Ribonucleic acids (RNAs) are versatile macromolecules that not only serve as carriers of genetic information, but also act as essential regulators and structural components influencing a wide range of life processes^1, 2^. RNA can be categorized into two main types: protein-coding RNA and non-coding RNA (ncRNA)^3, 4^. Protein-coding RNA primarily refers to message RNA^5^ which mainly functions by encoding genetic information through codons. NcRNA doesn’t encode proteins, instead, it regulates gene expression. NcRNA includes microRNA (miRNA), long non-coding RNA (lncRNA) and so on. Short miRNAs govern post-transcriptional gene regulation, while longer lncRNAs contribute to various cellular activities, from chromatin remodeling to epigenetic control.

For RNAs, sequence determines structure and structure determines function^6, 7^. The RNA diverse functions depend on the ability of single-stranded RNA molecules to fold into diverse secondary and tertiary structures. Understanding the structure of RNA is crucial for enhancing our overall knowledge of cellular biology and developing RNA-based therapeutics. Experimental methods like nuclear magnetic resonance^8^, X-ray crystallography^9^, cryogenic electron microscopy^10^, icSHAPE (in vivo RNA secondary structure profiles)^11^ have been developed to study the structure and function of RNA, which are expensive and time-consuming. Nowadays, approaches for RNA structure and function prediction can be divided into three categories, including thermodynamics-based^12–18^, alignment-based^19, 20^ and deep learning-based methods^21–25^. However, Thermodynamics-based methods are limited by inaccurate thermodynamic parameters. Alignment-based methods perform poorly when dealing with RNA sequences lacking MSA information. Although deep learning-based models demonstrate enhanced prediction accuracy across various datasets, these models usually performed worse when confronted with previously unseen data, suggesting that there is a notable drawback concerning their generalizability.

Nowadays, the advancements in high throughput sequencing technology^26^ have produced a wealth of unlabeled data, which contain rich information about RNA structures and functions. Many BERT-style^27^ RNA language models trained on abundant RNA sequences have been reported, such as RNA-FM^28^, RNABERT^29^, RNA-MSM^30^, CodonBERT^31^ and UNI-RNA^32^. RNA-FM is a BERT-based RNA foundation model trained on 23 million unannotated RNA sequences, demonstrating applications in predicting both structural and functional properties. RNABERT is developed by incorporating the Structure Alignment Learning (SAL) task during pre-training for the tasks like RNA structural alignment and RNA family clustering. RNA-MSM, also known as a MSA transformer^33^ for RNA, is developed based on evolutionary information for the tasks like base pair prediction and solvent accessibility. CodonBERT is a BERT-based language model trained on 10 million mRNA coding sequences, adopting a similar encoding approach as DNABERT^34^, and extended its application to predict various mRNA properties. UNI-RNA is a BERT-based language model trained on 1 billion RNA sequences with the largest parameters to date.

RNA structure is pivotal in elucidating its functional roles, but the existing pre-trained RNA language models fail to adequately incorporate structural information when parsing the RNA language space. BERT-based models, like RNA-FM trained with only one-dimensional sequences as input, may not be effective in extracting RNA structural and functional features. This could explain why the embedding extracted by RNA-FM is inferior to one-hot in certain tasks^30^. Furthermore, zero-shot RNA secondary structure prediction task also reveals that RNA-FM’s attention map does not capture RNA structural features explicitly. The attention map of RNA-MSM can be used to capture RNA secondary structure features on a specific test set with the MSA information, but this method is time-consuming and the performance decreases when dealing with RNA sequences with limited homologous sequences. The performance of current RNA language models indicate that BERT-based models designed for text feature extraction are not inherently suitable for extracting RNA features. Thus, it is necessary to explore whether integrating biological information, such as RNA structural information, into the computational framework of the BERT-based model can increase the performance.

The self-attention mechanism used in the transformer^35^-based models such as Alphafold2^36^ and Uni-mol^37^ plays a pivotal role in the feature extraction. Alphafold2 imposed an evolutionary and structural information-rich bias on the attention, which is well-suited to the iterative refinement of the protein structure. Uni-mol enhanced the molecular representation by replacing the attention map bias with the pair-wise interactions calculated from atom coordinates. Therefore, it is possible to utilize the key knowledge of RNA biology such as structural information to extract comprehensive representation of RNA features.

In this study, we developed a pre-trained RNA language model called ERNIE-RNA based on the modified BERT architecture by introducing a base-pairing informed attention bias when calculating attention, which enhanced the characterization of the RNA structure and enabled more thorough and comprehensive RNA feature extraction. Without any fine-tuning, the ERNIE-RNA’s attention maps were observed to successfully capture RNA structural features in a zero-shot manner, achieving an f1-score as high as 0.55. Through fine-tuning on diverse downstream tasks related to RNA structures and functions, ERNIE-RNA has achieved State-of-the-Art (SOTA) performance across all tested tasks, suggesting that ERNIE-RNA can be used to capture comprehensive RNA structural and functional information.

## Results

### The architecture of ERNIE-RNA

To address the limitations of traditional RNA language models, we introduce a novel RNA pre-trained language model, named ERNIE-RNA, which incorporates structural information based on the modified BERT model. The architecture of ERNIE-RNA contains 12 transformer blocks and each block contains 12 attention heads. Every token in the input sequences is mapped to a 768-dimensional vector, resulting in 86 million parameters. We utilized the one-dimensional RNA sequence to compute a pair-wise position matrix to replace the bias of the first layer in ERNIE-RNA. From the second layer onward, the bias of each layer is determined by the attention map of the previous layer. Pair-wise position matrix calculation involves iterating through every pair of bases i and j (where i ≠ j) in the given sequence with a length L. If the base pair at positions i and j is AU, the corresponding elements (i, j) and (j, i) in the matrix are set to 2. If it is a CG pair, these elements are assigned to 3. For GU pairs, the matrix elements (i, j) and (j, i) are assigned to a hyperparameter α, which is initially equal to 0.8. Lastly, all elements along the main diagonal of the matrix are set to 0. We collected 34 million non-coding RNA dataset from the RNAcentral^38^ database. After refining the vocabulary and removing redundant sequences, the final dataset consisted of 20.4 million RNA sequences. Subsequently, the ERNIE-RNA model was pre-trained on this dataset, which enabled the representation for RNA semantic information, including the L×L×156 attention map feature and the 12×768×L token embedding feature.

### ERNIE-RNA learns functional and structural information through pre-training

The ERNIE-RNA can extract various information simultaneously from the input sequences through multiple attention heads, which generate attention maps for illustrating the attention weight or importance assigned to different parts of input sequences by an attention mechanism. To investigate whether the attention map can provide valuable structural information, we conducted a zero-shot RNA secondary structure prediction experiment on the benchmark dataset bpRNA-1m^39^. We directly utilized the attention map from ERNIE-RNA as a probability map representing base-pairing in the RNA secondary structure. We employed the f1-score to evaluate the ability of each attention map to capture RNA secondary structures. Certain attention maps from ERNIE-RNA without fine-tuning were activated to effectively capture RNA secondary structure information with an f1-score of 0.55 (Figure 2a), which outperforms several methods fine-tuned on the bpRNA-1m training set, such as RNAfold^12^, RNAstructure^14^ and E2Efold^40^ (Table 1). Interestingly, we observed an increased capability of the attention map to capture RNA structural information as the attention map gets closer to the output layer, which may be due to the seamless transfer of attention information benefited from the ERNIE-RNA architecture.

**Figure 1:**
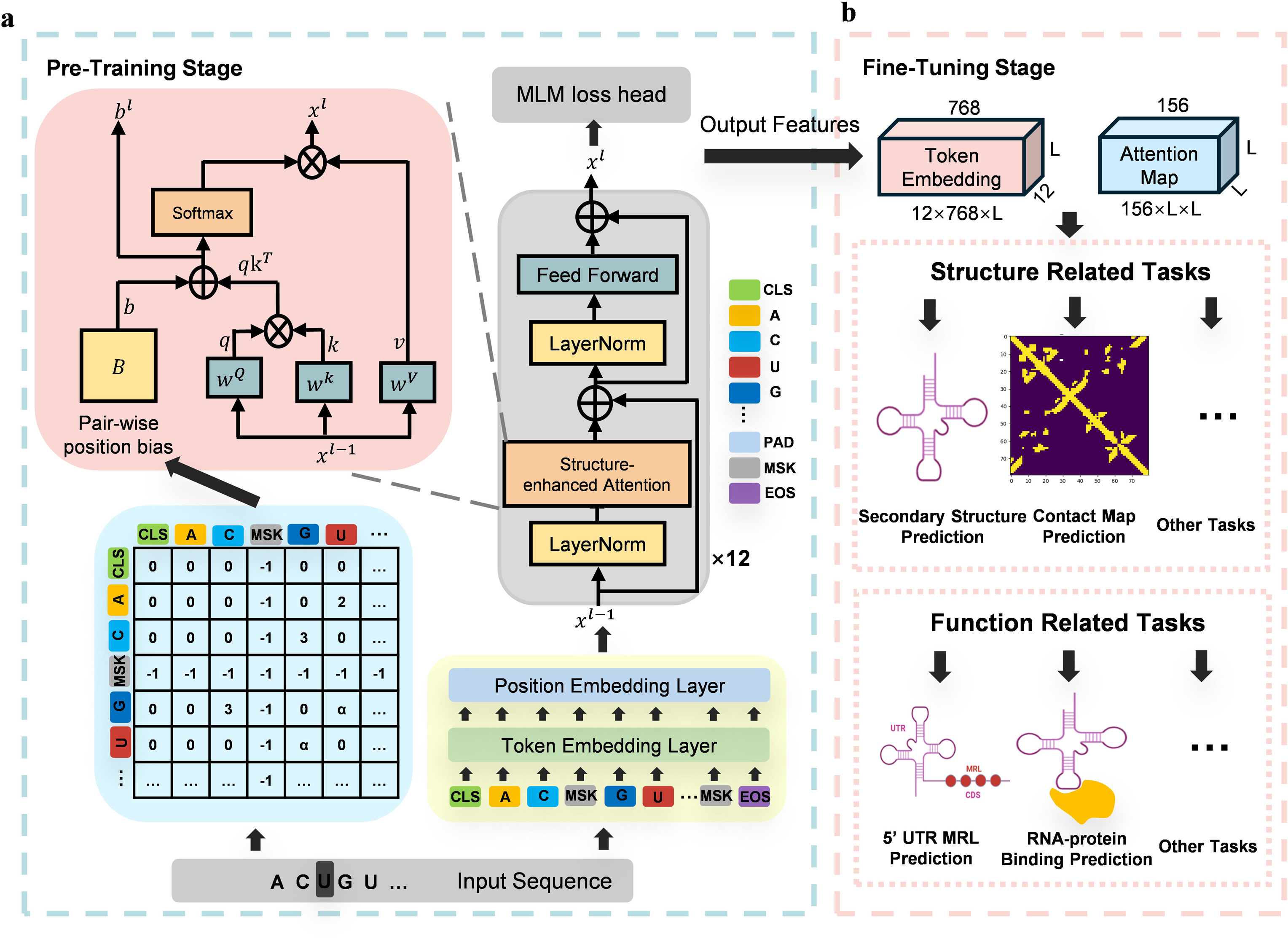
Overview of the ERNIE-RNA model architecture and application. ERNIE-RNA incorporates RNA structural information into the self-attention mechanism. **a**. In the pre-training stage, ERNIE-RNA consisting of 12 transformer layers was pre-trained with 20.4 million non-coding RNA sequences from RNAcentral via self-supervised learning. **b**. In the fine-tuning stage, ERNIE-RNA provides attention maps and token embeddings encoding rich structural and semantic RNA features, achieving state-of-the-art performance on diverse downstream tasks spanning structure prediction and functional annotation.

**Figure 2:**
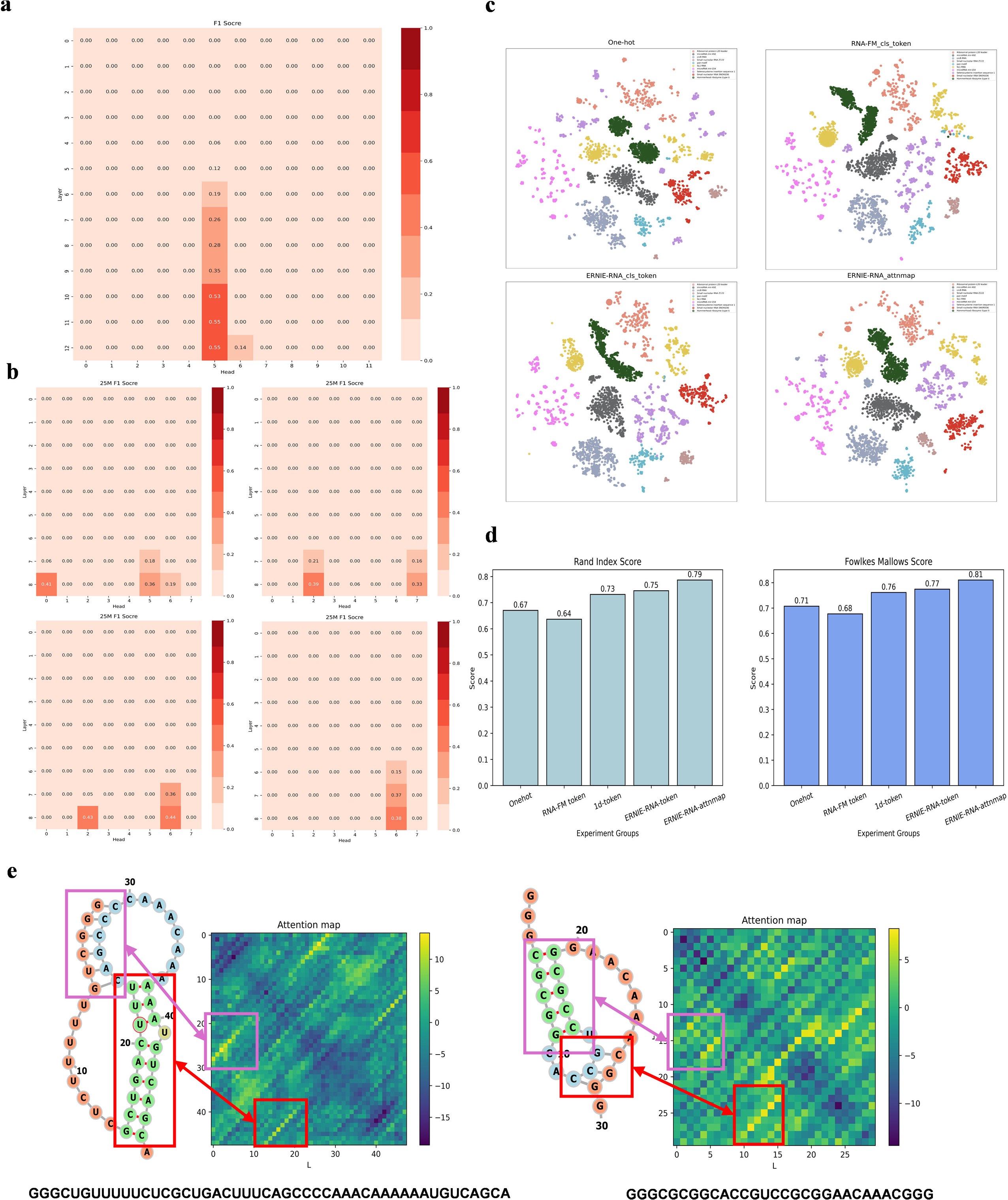
Zero-shot RNA secondary structure prediction experiment on bpRNA-1m test set and dimensionality reduction clustering experiment on small RNA dataset. **a.** Average binary f1 scores of 156 attention maps extracted from ERNIE-RNA (86M parameters; 12 attention heads; 12 transformer layers) on the bpRNA-1m test set. **b.** Average binary f1 scores of 72 attention maps from four 25M ERNIE-RNA with different initialization parameters (8 attention heads; 8 transformer layers) on the bpRNA-1m test set. **c.** Visualization of dimensionality reduction clustering results across features from 10 distinct small RNA categories. The best feature is attention map from ERNIE-RNA. **d.** Fowlkes-Mallows and Rand Index are used to quantitatively measure the disparities between cluster predictions and the true categories. **e.** The attention map extracted by ERNIE-RNA captures pseudoknot features on randomly selected two RNA sequences. 1-d base represents ERNIE-RNA without pair-wise position bias.

**Table1:**
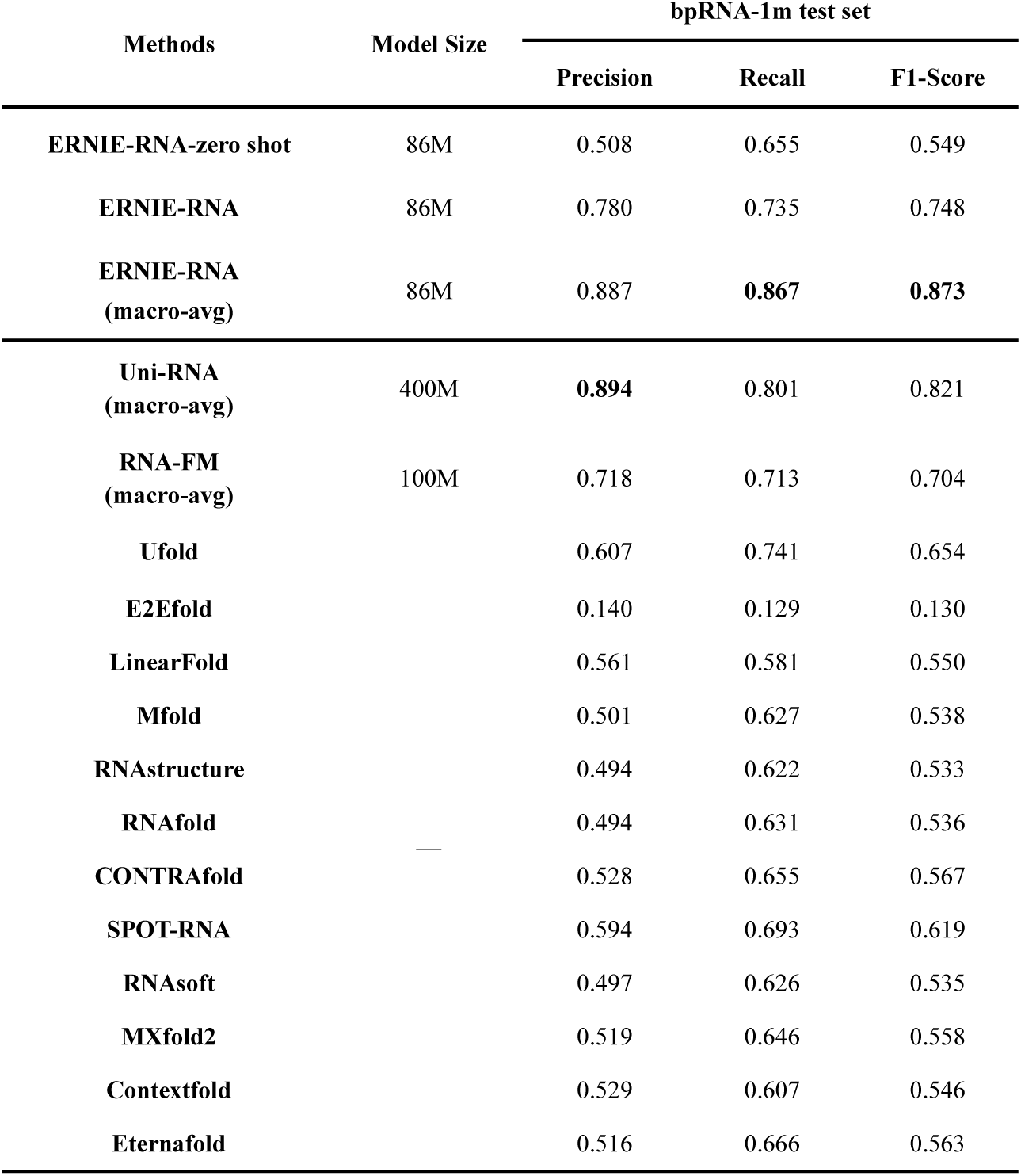
RNA secondary structure prediction performance. Despite not specifically designed for RNA secondary structure prediction, the ERNIE-RNA outperformed 14 other tested methods on all evaluation metrics. The benchmarks of other tested methods were adopted from RNA-FM and UNI-RNA paper. The (macro-avg) under several methods represents macro average f1-score, and other tested methods was evaluated with binary f1-score.

To investigate whether the presence of attention maps capturing RNA structural information is a coincidence, we further pre-trained four ERNIE-RNA models with 25 million parameters, initialized with different parameter values. In the similar zero-shot experiments, the capturing capability of RNA structural information was activated in different heads across different ERNIE-RNA models (Figure 2b). By comparison, no activated attention map was observed when a similar RNA model was pre-trained with no pair-wise position bias (supplementary Figure 1a). Furthermore, we did not observe any activated attention map in the ERNIE-RNA with random parameter values and without pre-training (supplementary Figure 1b). In addition, we performed the similar zero-shot experiment on the RNA-FM attention maps and none of the RNA-FM attention maps demonstrated the capability of capturing RNA secondary structure information (supplementary Figure 2).

Pseudoknot^41^ is a non-canonical type of secondary structure that can form in certain RNA molecules, characterized by base pairings between the loop region of a stem-loop structure and nucleotides outside that stem-loop. This knotted configuration makes pseudoknots challenging to accurately predict and analyze. Some traditional algorithms based on thermodynamics and dynamic programming, such as Mfold^42^, often omit pseudoknot prediction due to time complexity constraints^43^. We used ERNIE-RNA to extract attention maps from two RNA sequences containing pseudoknot motifs. As shown in Figure 2e, both attention maps successfully captured the sequences’ pseudoknot structural features, further demonstrating ERNIE-RNA’s capacity to learn RNA structural information after pre-training.

Token embedding extracted by ERNIE-RNA captures the semantic meaning of the individual token in the input RNA sequences and plays a crucial role in model’s ability to understand RNA language. We conducted a dimensionality reduction and clustering experiment on a small RNA dataset^44^. The dataset comprises 244 RNA families from RFAM^45^, including various RNA types such as tRNA, small nucleolar RNA and microRNA. To ensure the reliability of experiments, we performed ten rounds of repeated experiments. In each round, we randomly selected 10 RNA categories with lengths under 200 nucleotides and randomly sampled up to 1,000 sequences from each category. We extracted the CLS token embeddings of the RNA sequences by using RNA-FM and ERNIE-RNA with or without pair-wise position bias, and extracted attention maps only from ERNIE-RNA. We then employed t-SNE^46^ to reduce the feature dimensions to 2D and visualized the results. We found both CLS token embeddings and attention maps from ERNIE-RNA achieved superior performance compared to CLS token embeddings from RNA-FM, one-hot encoding features and ERNIE-RNA without pair-wise position bias, measured by using the Fowlkes-Mallows and Rand index (Figure 2c, 2d).

### ERNIE-RNA improves the performance of downstream tasks by fine-tuning on labeled data (Figure 3)

### RNA secondary structure prediction

We downloaded the benchmark dataset bpRNA-1m for RNA secondary structure prediction, which contains 13,419 RNA sequences with experimentally validated RNA structures. We randomly split the dataset into 10,814 training (TR0), 1,300 validation (VL0), and 1,305 testing (TS0) sets.

**Figure 3:**
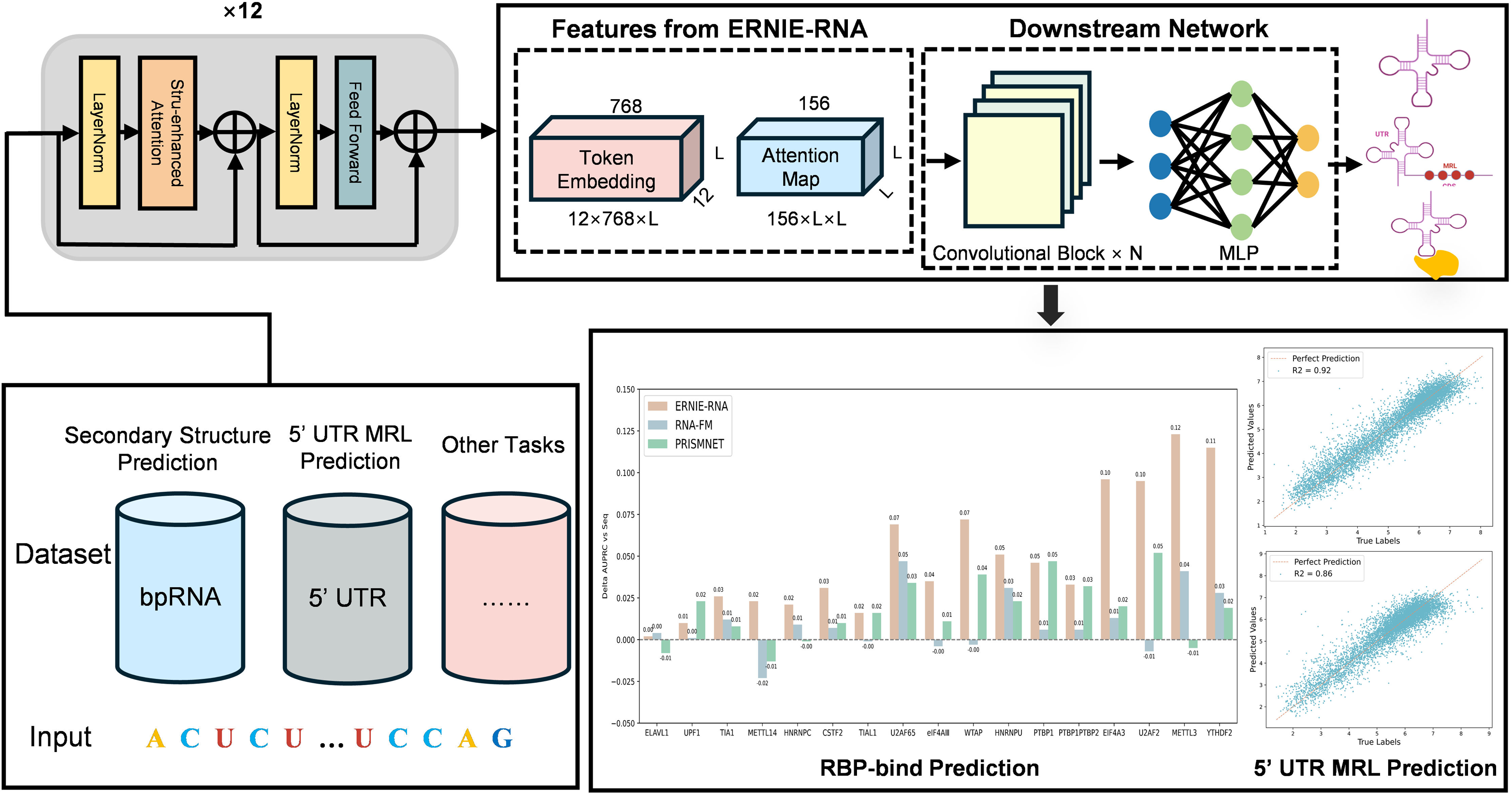
Framework diagram for downstream task fine-tuning based on ERNIE-RNA. Downstream network architectures comprise two broad categories: convolutional residual neural networks taking ERNIE-RNA’s attention maps or token embeddings as inputs, and fully connected feedforward networks using the CLS token embedding.

As shown in Table 1 and Figure 4a and 4c, ERNIE-RNA outperforms all other tested methods for RNA secondary structure prediction on the bpRNA-1m test set. Compared to UFold^22^ that is considered as the current state-of-the-art end-to-end method for RNA secondary structure prediction, ERNIE-RNA improves the average binary f1 score by 14.4% to 0.748. Despite ERNIE-RNA having less than one-fifth of the model size and pretrained with less than one-fiftieth of the data size compared to UNI-RNA^32^, ERNIE-RNA outperforms UNI-RNA, the currently largest pre-trained RNA language model, further boosting the average macro-average f1 score by 6.3% to 0.873. We further compared ERNIR-RNA’s prediction performance to RNA-FM at the single sample level. Each point represents one test sequence, as shown in Figure 4b. Clearly, The fined-tuned ERNIE-RNA provides better RNA secondary structure prediction results than RNA-FM in the majority of the samples. Among 1305 tested RNA sequences, binary f1-scores of 884 predicted by fine-tuned ERNIE-RNA are higher than that predicted by RNA-FM.

**Figure 4:**
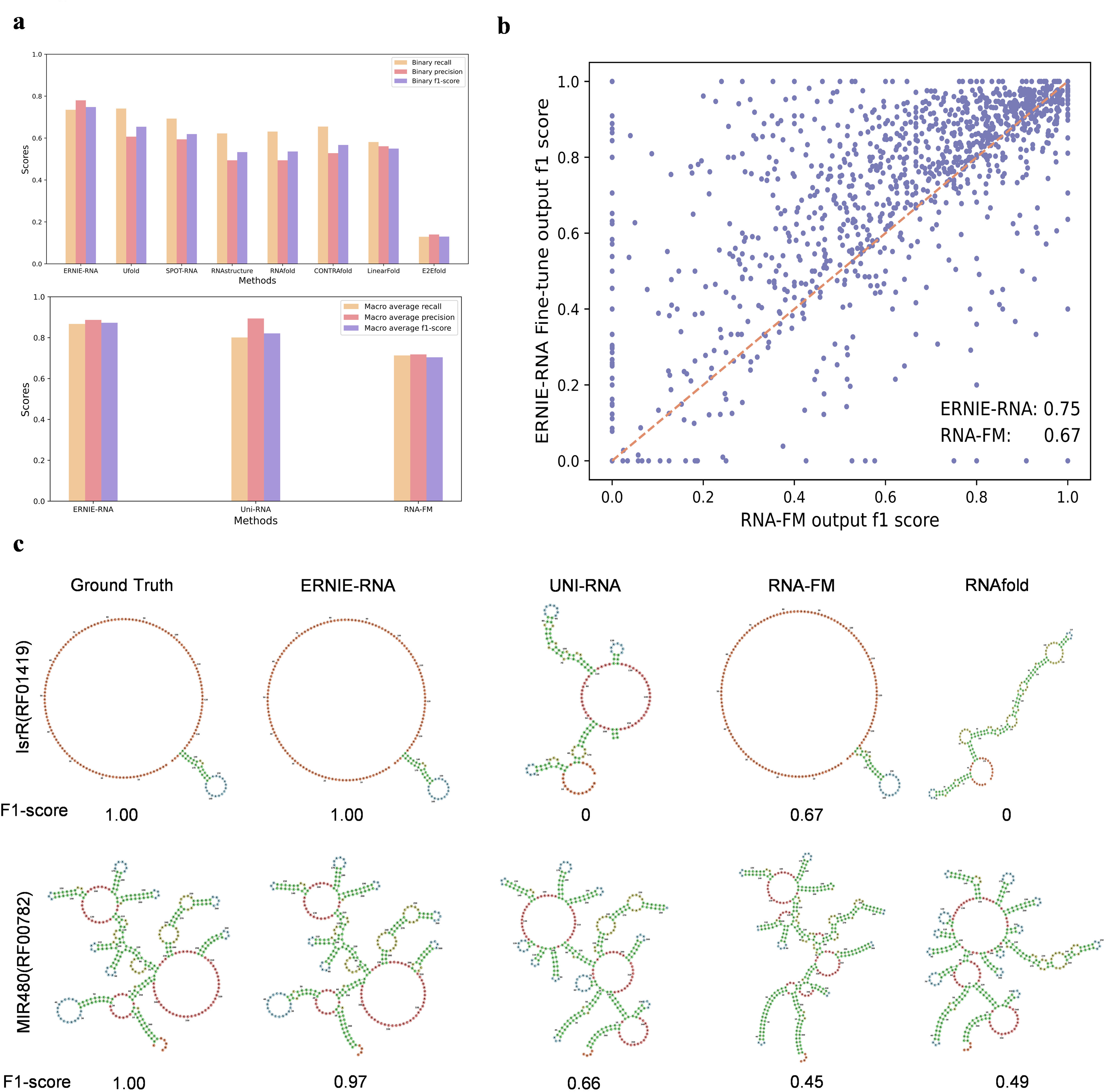
Performance of ERNIE-RNA on RNA structure prediction tasks. RNA secondary structure dataset and tertiary structure dataset: bpRNA-1m and PDB data**. a.** The bar chart compares the performance of different tested methods on the bpRNA-1m test set, with binary metrics (up) and macro average metrics (down). ERNIE-RNA outperforms all other tested methods on all kinds of f1-scores. **b.** The scatter plot benchmarks per-sample predictions of ERNIE-RNA against RNA-FM, with RNA-FM f1 scores on the x-axis and ERNIE-RNA F1 scores on the y-axis. **c.** RNA secondary structure predicted by different methods for two randomly selected samples. The first column is true label for each sample and f1-score represents binary f1-score.

### RNA contact map prediction

RNA contact map prediction task refers to predicting the spatial distances between nucleotides within RNA molecules based on their one-dimensional sequence. A RNA contact map employs a two-dimensional matrix wherein each cell indicates whether the distance between nucleotides at corresponding positions in the RNA molecule’s three-dimensional structure is below a predefined threshold (typically 8Å). Nucleotides falling within this threshold are in closer spatial proximity. We downloaded the benchmark datasets from RNAcontact^47^ that contains 301 sequences with >5 contacts, and we divided the dataset into 221 training (TR221) and 80 testing (TS80) sets.

We designed various models based on the ResNet downstream architecture but with different combinations of features, including one-hot, MSA (named Cov), RNA secondary structure predicted by PETfold^20^ (named SS), attention maps extracted from ERNIE-RNA and token embedding extracted from ERNIE-RNA and RNA-FM. Notably, the ResNet using attention maps from ERNIE-RNA as input outperformed all other tested models except for RNAcontact (ensemble), demonstrating 43% higher Top-L/1 precision compared to Cov+SS and 100% improvement over one-hot encodings (Figure 5a and Table 2). Although the RNAcontact (ensemble) showed a better performance, the ensemble output of 100 models used in RNAcontact (ensemble) may cause over-fitted results. These results indicated that the superior performance of ERNIE-RNA may be due to RNA structural features effectively encoded by the pre-training with the pair-wise position bias.

**Figure 5:**
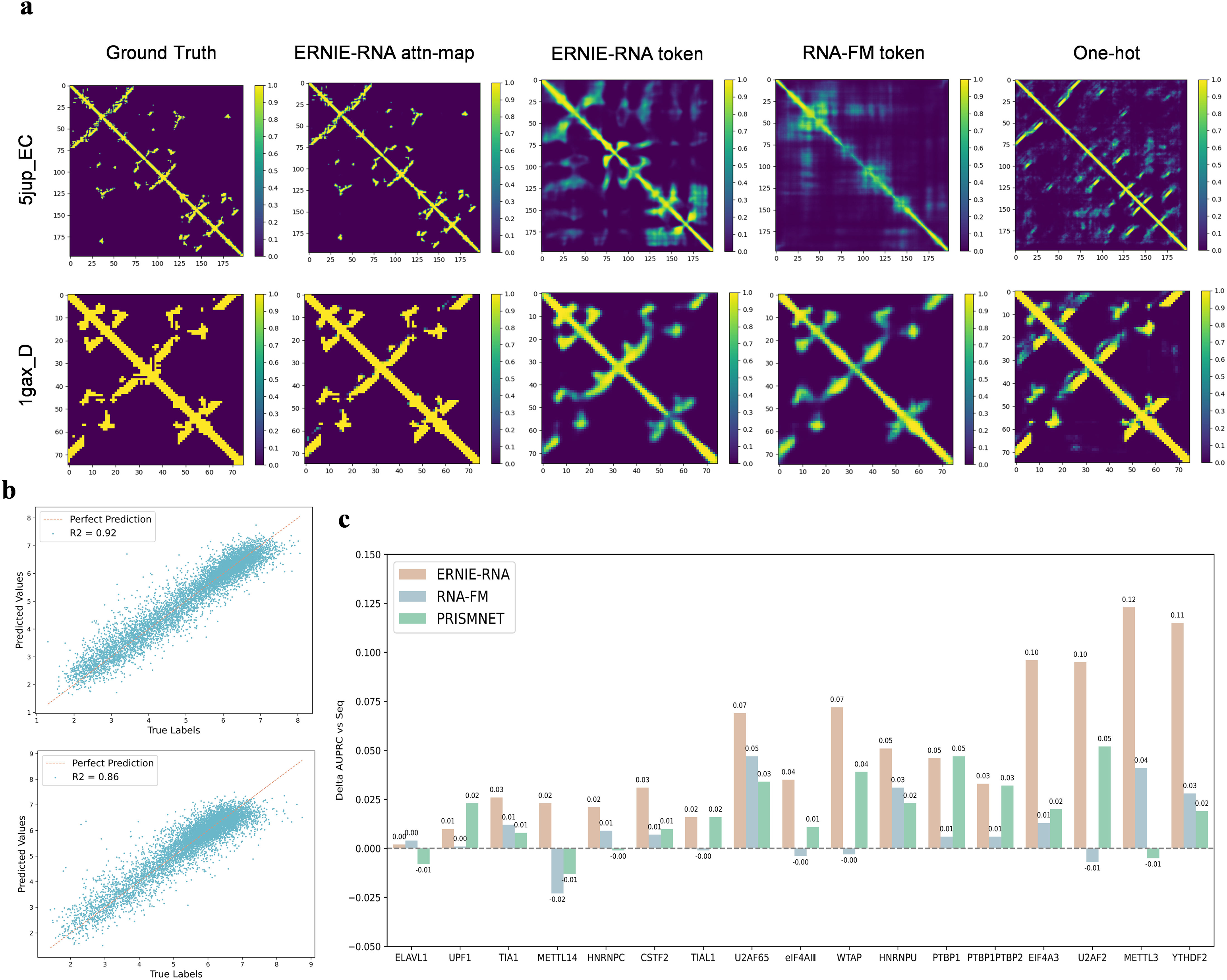
Performance of ERNIE-RNA on other downstream tasks. **a.** RNA contact maps of two randomly selected samples which are predicted by different methods. The first column is true label. **b**. ERNIE-RNA-conv’s performance (R²) on 5’UTR sequence MRL prediction dataset, including random test set (up) and human test set (down). **c**. Histogram depicting AUPRC distributions across 17 proteins compared to Seq baseline, denoted by the horizontal dashed line. ERNIE-RNA outperforms all other tested methods in most cases.

**Table2:** RNA contact map prediction performance. The first column of the table represents different features of RNA, ResNet represents the downstream network. The calculation methodology for the Long-Range Top Precision metric is detailed in the Methods section. The model using ERNIE-RNA’s attention map features outperformed all other tested methods except for RNAcontact with Cov+SS as input. The results of models that using one-hot or cov features as input were obtained from RNA-FM.

| Features | Methods | Long-Range Top Precision |  |  |  |
| --- | --- | --- | --- | --- | --- |
|  |  | L/10 | L/5 | L/2 | L/1 |
| One-hot | RNAcontact<br>(100 ensemble) | 0.48 | 0.45 | 0.40 | 0.33 |
| Token-Embedding<br>(RNA-FM) | ResNet | 0.56 | 0.55 | 0.49 | 0.42 |
| Cov | ResNet | 0.57 | 0.54 | 0.45 | 0.34 |
| Cov+SS | ResNet | 0.62 | 0.61 | 0.54 | 0.46 |
| Token-Embedding<br>(ERNIE-RNA) | ResNet | 0.65 | 0.62 | 0.56 | 0.47 |
| Attention-Map<br>(ERNIE-RNA) | ResNet | 0.81 | 0.78 | 0.74 | **0.66** |
| Cov+SS | RNAcontact<br>(100 ensemble) | **0.89** | **0.88** | **0.81** | 0.66 |

Cov represents multiple sequence alignment covariance features;
SS represents RNA secondary structure predicted by PETfold which use MSA information;
Token-Embedding represents all the token embeddings except for CLS token embedding and EOS token embedding;
+ represents channel-wise concatenation integrating different input representations.

### 5’UTR sequence MRL prediction

The 5’UTR (untranslated region) sequences MRL prediction task refers to predicting the mean ribosomal loading onto the 5’UTR sequences, which is often used to evaluate the translation efficiency of the corresponding RNA sequences. We downloaded the benchmark dataset from Optimus 5-prime^48^, which comprises 83,919 artificially synthesized random 5’UTRs and 7600 real human 5’UTRs along with corresponding mean ribosomal loading (MRL) values.

We selected 7,600 synthesized random 5’UTRs as the random test set with the remaining 76,319 synthesized random 5’UTRs sequences as the training set. The 7600 human 5’UTRs was used as the human test set to assess the model’s ability to generalize beyond synthetic 5’UTR sequences. We found that the performance of all tested models was worse on the human test set compared to the random test set. This may be due to distribution differences between the two datasets. As shown in Table 3 and Figure 5b, ERNIE-RNA-conv achieved the best performance on the random test set (R^2^=0.92) and the human test set (R²=0.86), which used all ERNIE-RNA’s token embeddings except for CLS and EOS token as input and convolutional residual neural network as the downstream architecture. Although ERNIE-RNA-mlp utilized only two simple mlp layers as its downstream architecture, fine-tuning performance is close to the SOTA (R^2^ =0.91 on the random test set and R²=0.84 on the human test set). Despite having the smallest model size and least pre-training data, ERNIE-RNA demonstrated the best generalizability for 5’UTR sequences MRL prediction task among all tested language models.

**Table3:** 5’UTR sequence MRL prediction performance (R^2^). We replaced the original input features of Optimus 5-Prime with token embeddings extracted from various language models. All pre-trained language models had better performance than traditional methods. Remarkably, ERNIE-RNA-conv outperfor

| Methods | Model Size | Random 7600 | Human 7600 |
| --- | --- | --- | --- |
| | | $R^2$ | |
| ERNIE-RNA-conv | 86M | **0.92** | **0.86** |
| ERNIE-RNA-mlp |  | 0.91 | 0.84 |
| UNI-RNA-conv | 400M | 0.91 | 0.85 |
| RNA-FM-conv | 100M | 0.85 | 0.79 |
| Optimus 5-Prime | - | 0.84 | 0.78 |
| FramePool |  | 0.82 | 0.78 |

-conv represents Convolutional Residual Neural Network, of which input is the whole token embedding except for CLS token and EOS token;
-mlp represents MultiLayer Perceptron, of which input is only CLS token embedding

### RNA-protein binding prediction

RNA-protein binding is a common biological phenomenon within cells and plays a very important role in various cellular activities, including cell-signaling and translation. We conducted experiments on the benchmark dataset from PrismNet^49^, which included icSHAPE data. We divided the dataset into several sub-datasets according to different corresponding RBPs (RNA-binding proteins) and different cell environment. We finally chose 17 RBPs in HeLa cell environment. We designed two models incorporating different feature representations extracted by ERNIE-RNA. All the models except for ERNIE-RNA (MLP) used the PrismNet as the same downstream network, only with different input features, including the CLS token embedding and the token embeddings without the CLS token.

As shown in Table 4 and Figure 5c, the model using icSHAPE and one-hot encoding features as input has a higher mean AUPRCs than that only with one-hot encoding features, which may be due to the RNA secondary structure information provided by icSHAPE. Notably, the ERNIE-RNA (MLP) which only use CLS token embedding as input performed better than all previous methods. Furthermore, the model replacing icSHAPE with token embeddings extracted by ERNIE-RNA is the best among all tested models, suggesting that ERNIE-RNA can learn sufficient information about structures and functions from raw RNA sequences, which benefits the downstream functional prediction task.

**Table4:**
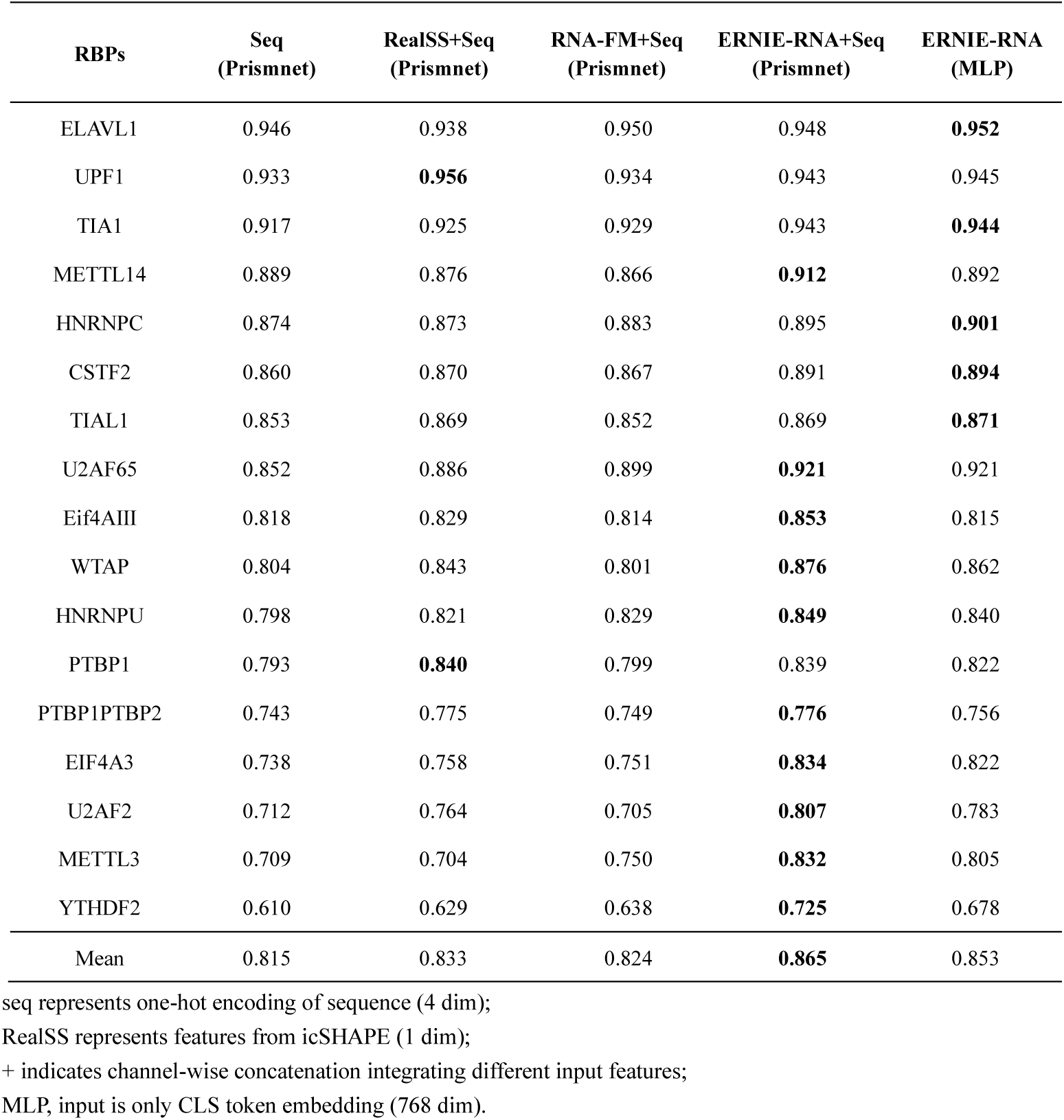
RBP-RNA binding prediction performance (AUPRC). Binary classification models were trained for each protein and mean ARPRCs was used to evaluate model performance. ERNIE-RNA+Seq and ERNIE-RNA (MLP) outperformed all other models except for UPF1 and PTBP1. ERNIE-RNA+Seq had the highest mean AUPRCs than others. Even ERNIE-RNA (MLP) was better than RealSS+Seq which used experimental data.

## Discussions

To effectively utilize the vast amount of unlabeled RNA sequences and extract RNA features with more comprehensive semantic information, we trained an RNA language model named ERNIE-RNA using 20.4 million non-coding RNAs from RNAcentral. Our results demonstrate that ERNIE-RNA’s attention maps inherently capture RNA structural features by pre-training alone. Upon fine-tuning, ERNIE-RNA achieves SOTA performance in downstream tasks like RNA secondary structure, RNA contact map, UTR-MRL and RNA-protein binding prediction. While ERNIE-RNA has offered some level of interpretability about RNA language models, deeper insights remain to be explored through further investigation. Furthermore, it is possible that when the ERNIE-RNA is pre-trained with limited RNA sequence data, simply increasing the number of model parameters may not lead to significant performance gains. Therefore, it is necessary to gather larger RNA sequence datasets to further investigate ERNIE-RNA’s potential with increased model size, which may result in emergent abilities of RNA language models. In summary, our results indicate that ERNIE-RNA may play a pivotal role in uncovering the fundamental principles governing the RNA biology, and the enhanced self-attention mechanism inspired by biological principles may also be used in pre-trained language models for proteins, DNA, and other biological molecules.

## Methods

### Training Dataset

We collected 34 million raw non-coding RNA (ncRNA) dataset from RNAcentral database, which is the largest dataset of ncRNA to date. We substituted T with U within the sequences and used 11 different symbols, namely ’N,’ ’Y,’ ’R,’ ’S,’ ’K,’ ’W,’ ’M,’ ’D,’ ’H,’ ’V,’ and ’B,’ to represent distinct degenerate bases, as illustrated in Table 5. After refining the vocabulary, CD-HIT-EST^50^ was used to remove redundant sequences above 100% similarity, resulting in 25 million non-redundant sequences. We further filtered sequences longer than 1024 and obtained a large-scale pre-training dataset consisting of 20.4 million ncRNA sequences finally. Figure 6 shows the length distribution and type distribution of this dataset.

**Figure 6:**
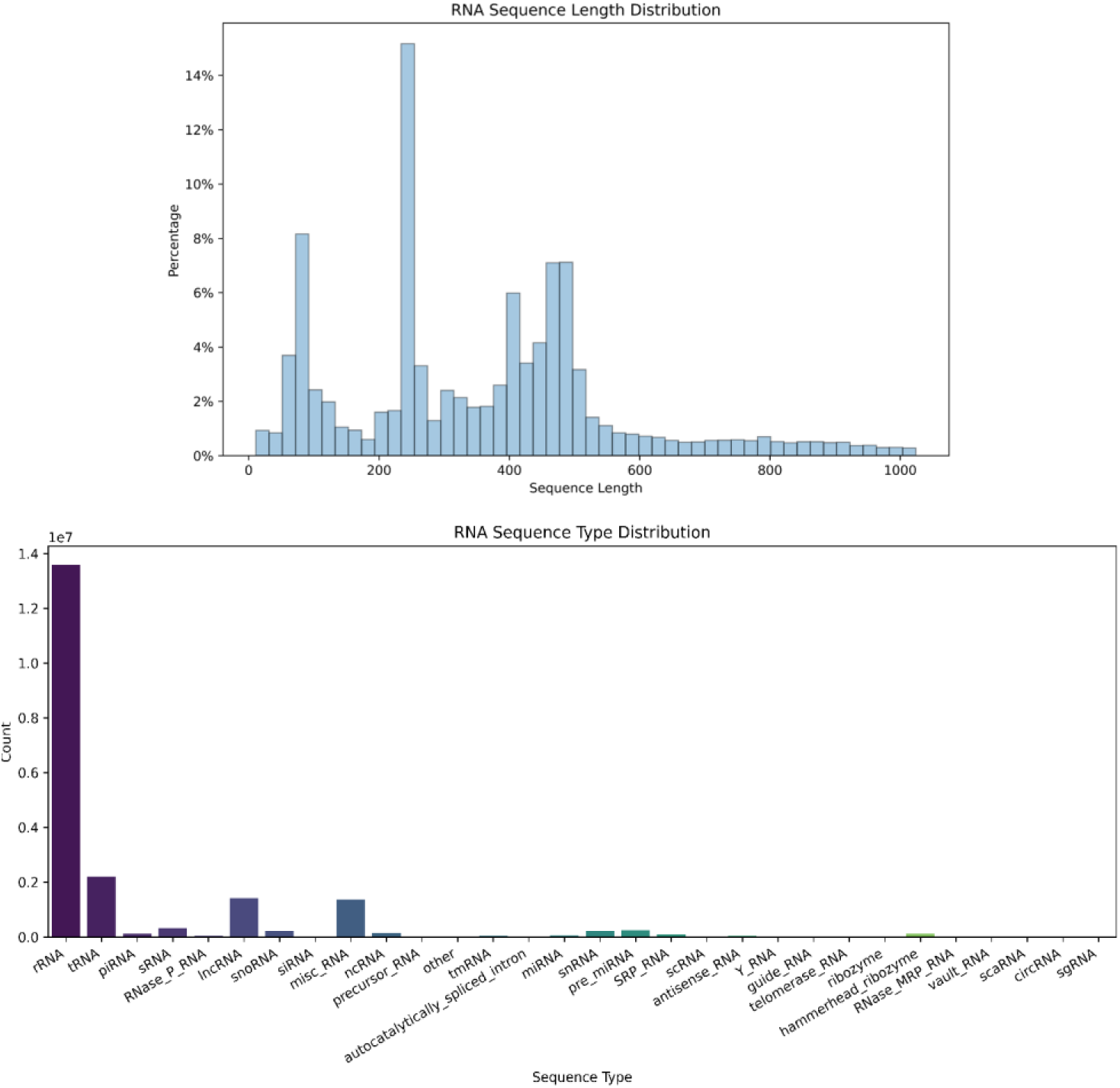
Length distribution and type distribution of 20.4M pre-training dataset.

**Table5:**
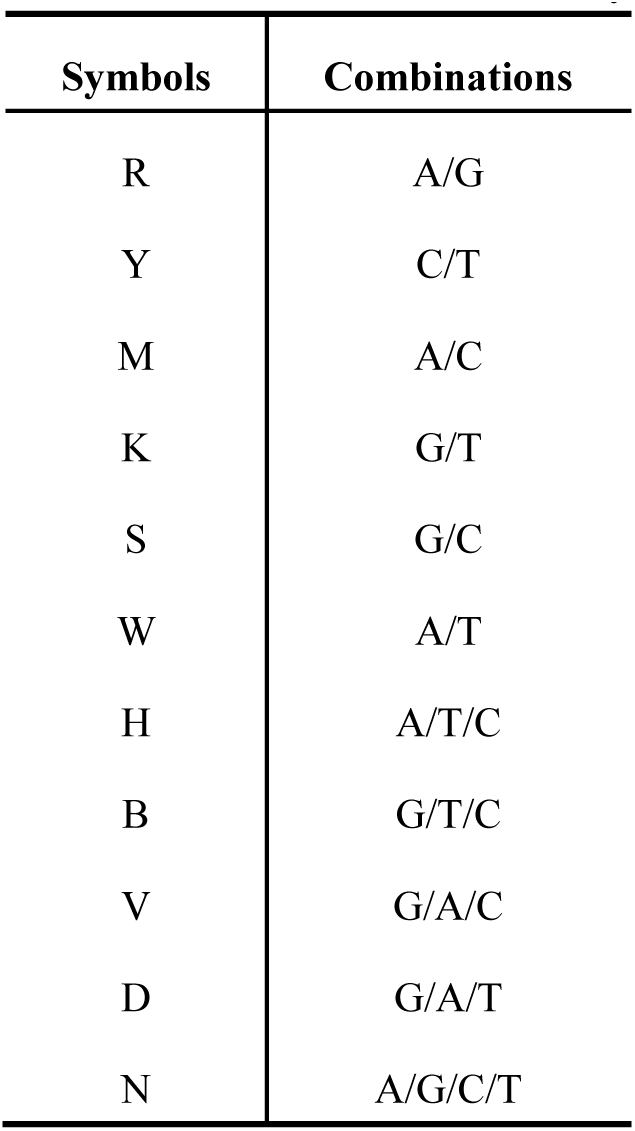
Base combinations of different symbols.

| Symbols | Combinations |
| --- | --- |
| R | A/G |
| Y | C/T |
| M | A/C |
| K | G/T |
| S | G/C |
| W | A/T |
| H | A/T/C |
| B | G/T/C |
| V | G/A/C |
| D | G/A/T |
| N | A/G/C/T |

### Model architecture

In this work, we introduced a RNA pre-trained language model, named ERNIE-RNA, which enhances structural information based on the modified BERT. ERNIE-RNA consists of 12 transformer blocks and each block contains 12 attention heads. Every token in the sequence is mapped to a 768-dimensional vector, resulting in 86 million parameters. Specifically, we utilized the one-dimensional RNA sequence to compute a pair-wise position matrix to replace the bias of the first layer in the ERNIE-RNA. From the second layer onward, the bias of each layer is determined by the attention map of the previous layer. This integration introduces RNA structural information into the attention map calculation at each layer, allowing for the extraction of more comprehensive semantic features. The improved self-attention formula is as follows:

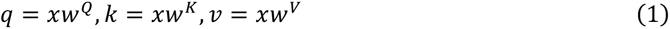

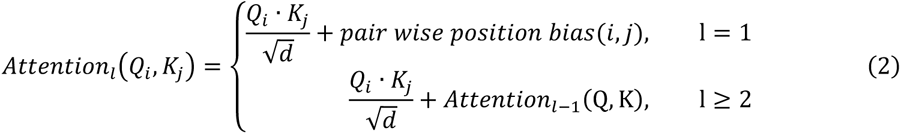

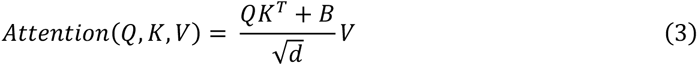

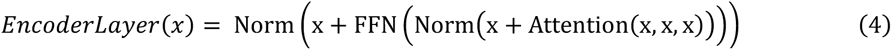

For a RNA sequence with length L, ERNIE-RNA takes raw sequential tokens as input, resulting in an L×768×12 embedding matrix and an L×L×156 attention maps, where 156 equals 12×13, 12 represents the num of attention heads and 13 represents the num of transformer blocks plus 1 (manually calculated pair-wise position matrix).

### Training details

For pre-training, we followed a self-supervised training manner in RNA-FM. Around 15% of nucleotide tokens are randomly replaced with a special token. (If the i-th token is chosen, we replace the i-th token with (1) the [MASK] token 80% of the time (2) a random token 10% of the time (3) the unchanged i-th token 10% of the time). We train ERNIE-RNA with masked language modeling (MLM), which predict the masked token with cross-entropy loss.

We use fairseq^51^ to train ERNIE-RNA for about 20 days on 24 32G-V100. During the pre-training process, we adopted the following hyperparameter configuration: the base learning rate was set to 0.0001, the warmup step was set to 20000 steps, and the weight-decay was set to 0.01. In order to speed up the training of the network while reducing the memory usage, we constrain the maximum length of the input sequence to 1024.

### Downstream dataset

#### RNA secondary structure dataset

The benchmark dataset bpRNA-1m is built according to Ufold and RNA-FM. This dataset was preprocessed by removing sequences with >80% identity and restricting maximum lengths under 500 nucleotides. The resulting dataset contains 13,419 sequences, randomly split into 10,814 training (TR0), 1,300 validation (VL0), and 1,305 testing (TS0) sets. Our data splitting ensures consistency with previous work to allow fair comparison. Different from UNI-RNA which use RNAstralign and TR0 as training dataset, we only use TR0 to train ERNIE-RNA and test on TS0.

#### RNA 3D closeness dataset

We utilized the benchmark datasets from Leontis and Zirbel. The original 1,786 sequences were downloaded from PDB and those lengths outside the range of 32 to 1000 were first excluded, leaving 511 sequences remained. CD-HIT-EST and BLASTclust^52^ were used to remove redundant sequences above 30% similarity, resulting in 336 non-redundant sequences. The distance between any two bases was defined as the minimum atom-pair distance. Bases’ distance within 8Å were labeled positive contacts. Finally, sequences with <5 contacts were removed, leaving 301 sequences divided into 221 training (TR221) and 80 testing (TS80) sets. In order to select the model checkpoint more fairly, we further randomly divided TR221 into two parts according to the ratio of 8:2, recorded as TR168 and VL43.

#### RNA 5’UTR mean ribosome loading dataset

Our experiments relied on a benchmark dataset obtained from Optimus 5-prime, including 83,919 artificially synthesized random 5’UTRs with their corresponding mean ribosomal loading (MRL) values. These sequences ranged from 25 to 100 nucleotides in length. To ensure the accuracy of our model testing across various 5’UTR lengths, we meticulously selected the top 100 5’UTRs of each length, prioritizing those with the deepest sequencing and highest read counts to enhance confidence in sequencing outcomes. This yielded a total of 7,600 sequences for the test set, while the remaining 76,319 sequences constituted the training set. Additionally, we enriched our analysis by incorporating an extra dataset comprising 7,600 real human 5’UTRs with the similar length distribution.

#### RNA protein binding dataset

We conduct experiments on benchmark dataset from PrismNet, which includes icSHAPE data. We divided them into several sub-datasets according to different corresponding RBPs and different cell environment. We finally chose 17 RBPs in HeLa cell environment as RNA-FM did to make a fair comparison. The number of RNA sequences of each RBP ranges from 1827 to 15002. We partitioned 20% of the data into a test set following an 8:2 ratio. Subsequently, the remaining 80% of the data was divided into a training set and a validation set at a ratio of 9:1. The length of all sequences from different RBPs is 101 nucleotides.

### Downstream tasks

#### RNA secondary structure & 3D closeness prediction

As both RNA secondary structure and the 3D contact map can be effectively represented by two-dimensional matrices, we used same downstream networks architecture in RNA secondary structure & 3D closeness prediction tasks. We designed the models based on ERNIE-RNA which is followed by several CNN layers. ERNIE-RNA uses pre-trained model parameters and the parameters of downstream CNN layers are initialized randomly. The models took token embedding and attention maps extracted by ERNIE-RNA as input. In the fine-tuning stage, we set the base learning rate to 1e-5 and batch size was set to 1; We use the cross-entropy loss function to update the model parameters. According to statistics based on bpRNA-1m, we set the weight ratio of loss function to 300:1; We also used an early stopping mechanism during the training process, when the performance of validation set does not improve for 20 consecutive epochs, model training will be terminated early. For RNA secondary structure prediction task, we used the same postprocess as E2Efold.

The Long-Range Top-L precision is a metric used to evaluate the accuracy of long-range contact prediction in the RNA contact map prediction task. The “Long-Range” aspect focuses on evaluating the effectiveness of base contact prediction for nucleotides with a distance greater than 24 (|i-j|≥24) in the contact map. So the predicted contact map’s elements with a difference in their x and y coordinates (|i-j|) less than or equal to 24 are set to 0. For example, the computation process for L/10 precision involves sorting all the values in the contact map from highest to lowest and selecting the top L/10 values as the predicted positive samples. The precision is then calculated based on the proportion of true positive samples among these selected samples. The Long-Range Top Precision is further divided into four categories: L/10, L/5, L/2, and L/1, all ranging between 0 and 1, with a higher value indicating better model performance.

#### RNA 5’UTR mean ribosome loading prediction

Prior to model construction and training, we firstly standardized the Mean Ribosomal Loading (MRL) values to be predicted, which may improve the convergence and overall performance of the model during fine-tuning. Two distinct models were constructed: ERNIE-RNA-token-conv and ERNIE-RNA-token-mlp. The former used Token Embedding except for CLS and EOS token from ERNIE-RNA, and employed a convolutional residual neural network as its downstream architecture. Conversely, ERNIE-RNA-token-mlp utilized the CLS Token Embedding provided by ERNIE-RNA to mitigate the impact of sequences of varying lengths. It employed a simple two-layer fully connected network to further extract features. During the fine-tuning phase, we initialized the base learning rate to 1e-5 and employed the Mean Squared Error (MSE) loss function to iteratively update model parameters. The number of tolerable epochs is 10. Considering that the design of ERNIE-RNA-token-conv is related to the input length, and the longest sequence length in the training set is 100, we applied padding to standardize the sequence length across the entire dataset to 100 for model input.

#### RNA protein binding prediction

We designed two models: ERNIE-RNA-PRSIMNET and ERNIE-RNA-MLP. ERNIE-RNA-PRSIMNET adopted the main architecture of PrismNet, but replaced the icSHAPE input with token embedding except for CLS and EOS token from ERNIE-RNA. ERNIE-RNA-MLP only took the CLS token embedding provided by ERNIE-RNA as input and further extracted features through a straightforward two-layer fully connected network. During the fine-tuning phase, we initialized the base learning rate to 1e-5 and employed the cross-entropy loss function to update model parameters. Additionally, we set 10 as tolerable epochs.

## Code availability

The scripts are available at https://github.com/Bruce-ywj/ERNIE-RNA

## Acknowledgements

We thank members of Xie lab for helpful discussions. This research is supported by the National Key Research and Development Program of China (2021YFC2302401 to G.L.; 2019YFA0906103 to Z.X.), National Natural Science Foundation of China (No. 32341014 [Z.X.]), Beijing Natural Science Foundation (No. Z230015 [Z.X.]).

## Author Contributions

Z.X. and W.Y. conceived of the ideas implemented in this project. W.Y. and L.H. designed the ERNIE-RNA models. W.Y. performed the pre-trained and downstream experiments, S.Z. assisted with experiments. Z.Z., X.Z., T.Q. and L.H. analyzed the results, R.J. and G.L. assisted with analysis. Z.X. supervised the project. Z.X. and W.Y. wrote the paper.

