## Supplementary Figure 1 for "ERNIE-RNA: An RNA Language Model with Structure-enhanced Representations"

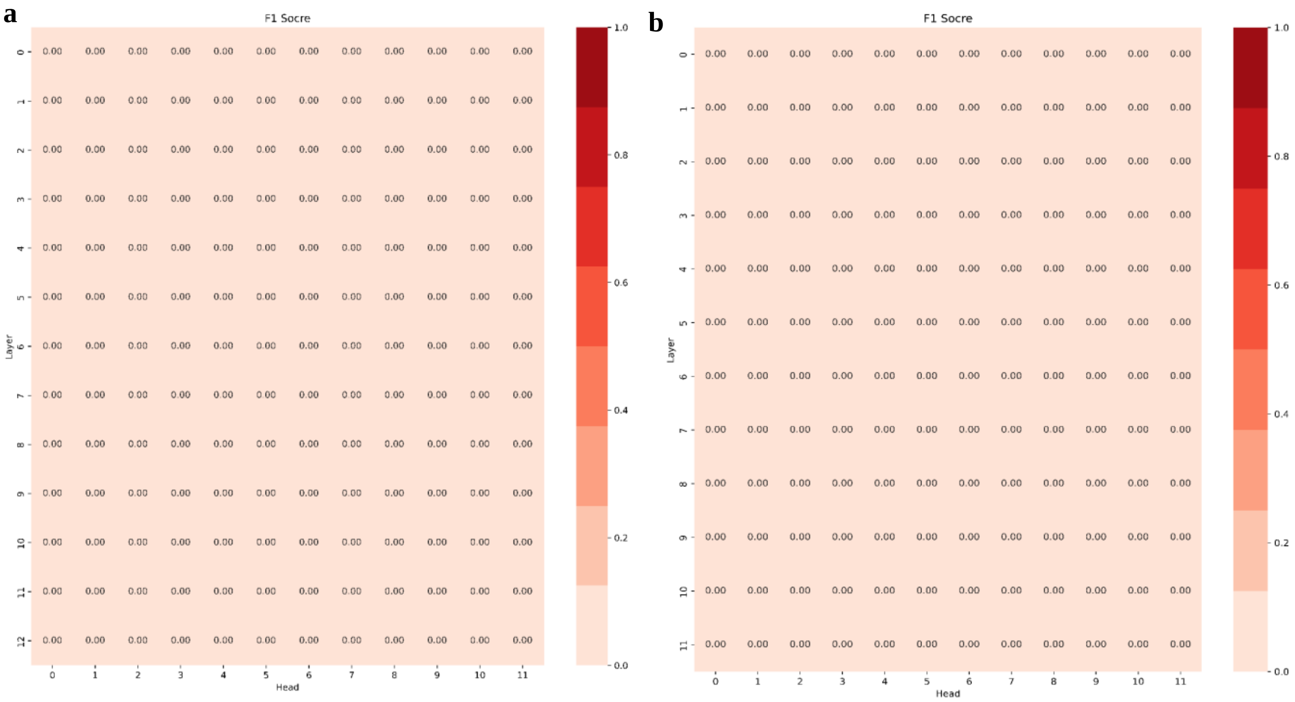


**Supplementary Figure 1: Zero-shot RNA secondary structure prediction experiment. a.** 1-d base model which has the exact same parameter configuration as ERNIE-RNA, except for the pair-wise position bias. **b.** ERNIE-RNA with random parameters. None of attention maps from above two models demonstrated capability in capturing RNA secondary structure information, all f1-scores are 0.

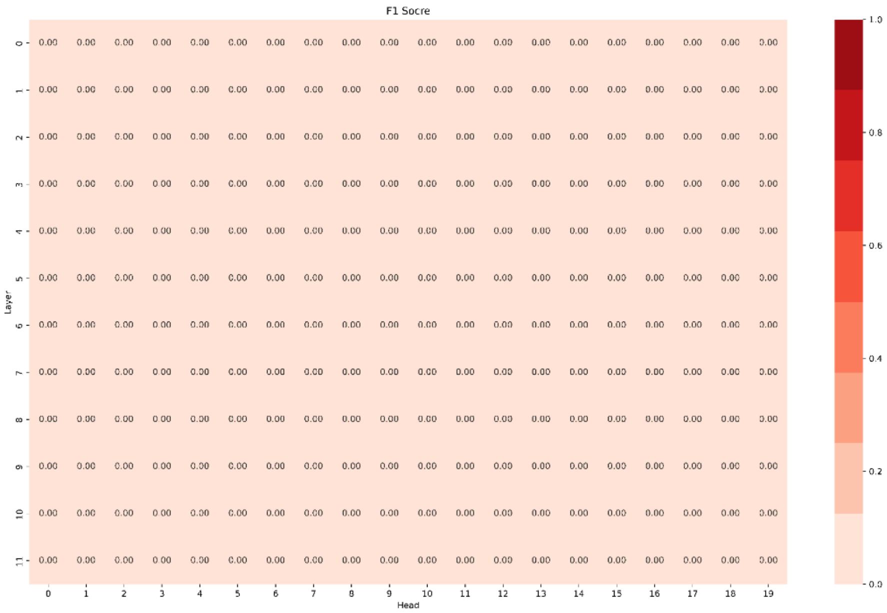


**Supplementary Figure 2: Zero-shot RNA secondary structure prediction experiment on RNA-FM.** None of the RNA-FM attention maps demonstrated capability in capturing RNA secondary structure information, all f1-scores are 0.
